# Two Nimrod receptors, NimC1 and Eater, synergistically contribute to bacterial phagocytosis in *Drosophila melanogaster*

**DOI:** 10.1101/479550

**Authors:** Claudia Melcarne, Elodie Ramond, Jan Dudzic, Andrew J Bretscher, Éva Kurucz, István Andó, Bruno Lemaitre

## Abstract

Eater and NimC1 are transmembrane receptors of the *Drosophila* Nimrod family, specifically expressed in hemocytes, the insect blood cells. Previous *ex vivo* and *in vivo* RNAi studies have pointed to their role in the phagocytosis of bacteria. Here, we have created a novel null mutant in NimC1 to re-evaluate the role of NimC1, alone or in combination with Eater, in the cellular immune response. We show that NimC1 functions as an adhesion molecule *ex vivo*, but in contrast to Eater is not required for hemocyte sessility *in vivo*. *Ex vivo* phagocytosis assays and electron microscopy experiments confirmed that Eater is the main phagocytic receptor for Gram-positive, but not Gram-negative bacteria, and contributes to microbe tethering to hemocytes. Surprisingly, the *NimC1* deletion did not impair phagocytosis of bacteria, nor their adhesion to the hemocytes. However, phagocytosis of both types of bacteria was almost abolished in *NimC1^1^*;*eater^1^* hemocytes. This indicates that both receptors contribute synergistically to the phagocytosis of bacteria, but that Eater can bypass the requirement for NimC1. Finally, we uncovered that NimC1, but not Eater, is essential for uptake of latex beads and zymosan particles. We conclude that Eater and NimC1 are the two main receptors for phagocytosis of bacteria in *Drosophila,* and that each receptor likely plays distinct roles in microbial uptake.

## Introduction

Phagocytosis is an ancient and evolutionarily conserved process, generally defined as the cellular uptake of particles bigger than 0.5 µm. Phagocytosis is an important feeding mechanism in primitive and unicellular organisms, such as amoeba [1]. In higher organisms, phagocytosis is performed by dedicated cells, generally called phagocytes, and is necessary for a plethora of specialized biological functions. *In primis*, phagocytosis is used as a powerful process to internalize and eliminate pathogens, as well as to trigger host inflammation [2]. Moreover, phagocytosis contributes to tissue homeostasis and embryonic development, mainly via the removal of apoptotic corpses [3]. Phagocytosis is a complex membrane-driven process guided by the actin cytoskeleton of the host phagocytic cell. It involves the recognition and subsequent binding of the microbe by surface receptors. These interactions are essential to activate intracellular signalling pathways that finally culminate in the formation of the phagosome [4]. Several studies have highlighted similarities between the phagocytic machinery of *Drosophila* and mammals, such as the involvement of actin and actin-related proteins [5]–[7]. *D. melanogaster* harbours highly efficient phagocytes, called plasmatocytes, which originate from multipotent progenitors (prohemocytes). In healthy larvae, prohemocytes can differentiate into two mature hemocyte types: plasmatocytes and crystal cells, which are involved in melanization response [8]. Plasmatocytes are professional phagocytes sharing functional features with mammalian macrophages, and represent the most abundant hemocyte class at all developmental stages. They play a key role in bacterial clearance during infection, as well as in the removal of apoptotic corpses [9], [10]. In addition to their phagocytic tasks, plasmatocytes also produce antimicrobial peptides [11] and cytokines [12], and synthesize extracellular matrix components [13]. *Drosophila* haematopoiesis occurs in two successive waves. A first set of hemocytes is produced during embryogenesis, giving rise to a defined number of plasmatocytes and crystal cells. This embryonic hemocyte population expands in number during the following larval stages. The second hemocyte lineage derives from the lymph gland, a specialized organ that develops along all larval stages. The lymph gland acts as a reservoir of both prohemocytes and mature hemocytes, which are released at the onset of metamorphosis or upon parasitization [8], [14]–[17]. Both blood cell lineages persist into the adult stage of the fly [18]. In the *Drosophila* larva, in addition to the lymph gland and the hemolymph, hemocytes can also be found in a third compartment: the sessile hematopoietic tissues [19]–[23]. There, hemocytes attach to the inner layer of the cuticle, forming striped patterns along the dorsal vessel, and lateral patches in association with the endings of peripheral neurons [8], [20], [22], [24]. Accumulating evidence suggests that this sub-epidermal sessile compartment of hemocytes functions as an active peripheral hematopoietic niche [20], [22], [23].

The ability of *Drosophila* hemocytes to perform efficient phagocytosis, relies on the expression of specific cell surface receptors that can bind particles and induce their engulfment. While many receptors have been implicated in bacterial phagocytosis, their specific involvement or individual contribution is less clear [25], [26]. In this paper, we have characterized the phagocytic role of NimC1 and Eater, two EGF-like repeat Nimrod surface receptors specifically expressed in hemocytes (Kocks et al., 2005; Kurucz et al., 2007). The Nimrod family of proteins is characterized by the presence of epidermal growth factor (EGF)-like domains, also called “NIM repeats”. This family comprises a cluster of ten proteins (NimA, NimB1-5 and NimC1-4) encoded by genes clustered on the chromosome II, and two related hemocyte surface receptors, Eater and Draper, encoded by genes on chromosome 3 [27], [28]. Early studies have shown the implication of some Nimrod C-type proteins in bacterial phagocytosis (Eater and NimC1) [27], [29], [30] or engulfment of apoptotic bodies (NimC4 or SIMU) [31], [32]. More recently, the Eater transmembrane receptor has also been involved in hemocyte adhesion and sessility [30]. Nimrod C1 (NimC1) is a 90 kDa transmembrane protein characterized by 10 NIM repeats in its extracellular region, a single transmembrane domain and a short cytosolic tail with unknown function [27]. NimC1 has been initially identified as the antigen of a hemocyte-specific antibody (P1), being involved in phagocytosis of bacteria [27]. Kurucz and colleagues showed that *NimC1* silencing by RNAi decreases *Staphylococcus aureus* uptake by plasmatocytes, whereas its overexpression in S2 cells enhances phagocytosis of both *S. aureus* and *E. coli* bacteria and makes the cells highly adherent [27]. Here, we generated a null mutation in *NimC1* by homologous recombination (called *NimC1*^1^) and revisited its function in hemocyte-mediated immunity. Moreover, we recombined the *NimC1* mutation with the previously described *eater*^1^ mutant [30], generating a *NimC1^1^;eater^1^* double mutant. Using these genetic tools, we first show the involvement of NimC1 in *ex vivo* cell adhesion and in the regulation of hemocyte proliferation. Contrasting with previous RNAi studies [27], our *ex vivo* phagocytosis assays demonstrate that NimC1 is not required for phagocytosis of Gram-positive or Gram-negative bacteria. Nevertheless, we show that this Nimrod receptor contributes to the uptake of latex beads and zymosan yeast particles. The use of the *NimC1^1^;eater^1^* double mutant not only re-confirmed Eater as the main Gram-positive engulfing receptor, but, more importantly, revealed a synergistic action of NimC1 and Eater in microbe phagocytosis. *NimC1^1^;eater^1^* hemocytes from third instar larvae, failed indeed to phagocytose any type of bacteria. Collectively, our study points to a major role of NimC1 and Eater in the phagocytosis of bacteria, and suggests that those proteins likely play distinct roles in microbial uptake, as tethering and docking receptors.

## Materials and Methods

### *Drosophila* stocks and methodology

All *Drosophila* stocks were maintained at 25°C on standard fly medium consisting of 6% cornmeal, 6% yeast, 0.62% agar, 0.1% fruit juice, supplemented with 10.6 g/L moldex and 4.9 ml/L propionic acid. Second instar (L2) larvae were selected 48-52 hours after egg laying (AEL), middle L3 larvae 72-90 hours AEL and third instar (L3) wandering larvae 110-120 hours AEL.

Wild type *w*^*1118*^ (BL5905) flies were used as controls, unless indicated otherwise. The following fly lines were used in this study:

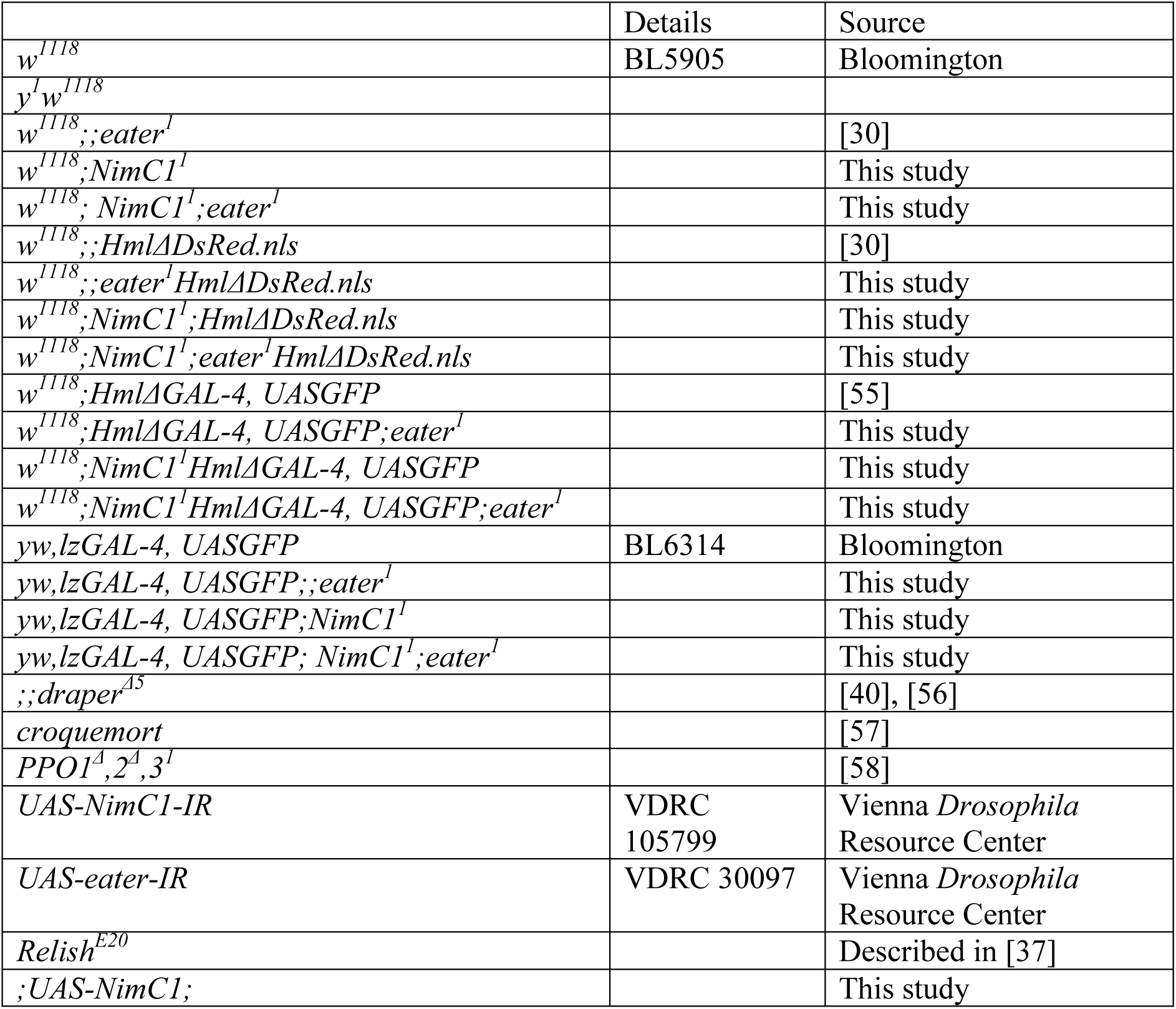

### Gene targeting of *NimC1*

Gene targeting of *NimC1* was performed as follows. The 5’ and 3’ homology arms, of 4.8kb and 3.7kb, respectively, were PCR amplified from genomic DNA. The 5’ arm was inserted between *NotI* and *NheI* restriction sites, whereas the 3’ arm was inserted between *SpeI* and *AscI* sites of the gene targeting vector pTV[Cherry]. A donor transgenic stock was generated by transformation of a starting *w1118 (*BL5905) stock, and used for hsFLP and hs-I-SceI mediated gene targeting [33]. Using this method, we recorded a 1/2000 knockout efficiency of the F2 progeny, i.e. 1/2000 offspring were bonafide *NimC1* knockouts.

The following primers were used for PCR genotyping and for testing the functional *NimC1* deletion by RT-PCR:

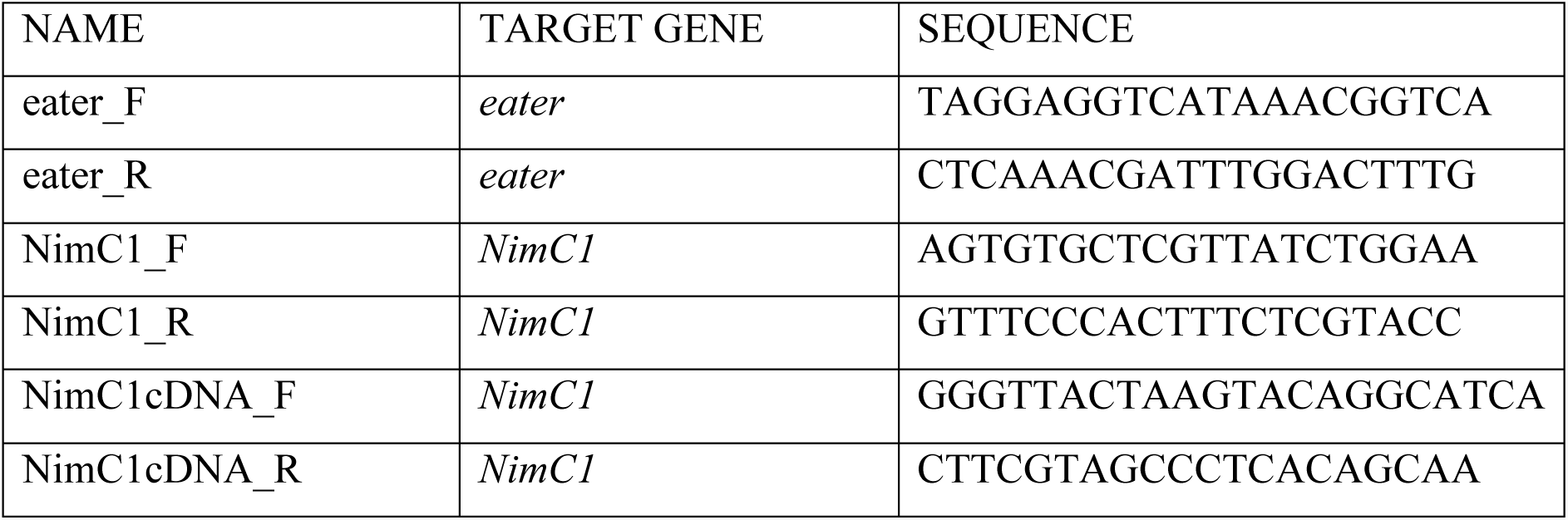

### Hemocyte counting by flow cytometry

A BD Accuri C6 flow cytometer was used to analyse hemocyte numbers. For each genotype, 15 L2, 5 middle L3, or 5 L3 wandering third instar larvae containing the *HmlΔdsred.nls* marker were bled into 150 µL of Schneider’s insect medium (Sigma-Aldrich) containing 1 nM phenylthiourea (PTU, Sigma), and 100 µL of hemocyte suspension was analysed by flow cytometry. Hemocytes were first selected from debris by plotting FSC-A against SSC-A on a logarithmic scale in a dot plot. Cells were then gated for singlets by plotting FSC-H versus FSC-A. *w*^*1118*^ and *w*^*1118*^;;*HmlΔdsred.nls* larvae were used to define the gates for hemocyte population, using a FL2 detector.

### Hemocyte size measurement of free floating cells

Third instar (L3) wandering larvae were bled in 1X PBS without calcium and magnesium, supplemented with EDTA 5mM. Invitrogen™ Tali™ Image-based Cytometer machine was used to measure hemocytes size in suspension of more than 7 000 cells per genotype.

### *Ex vivo* larval hemocyte phagocytosis assays

1. *Ex vivo* phagocytosis assay of *Escherichia coli* and *Staphylococcus aureus* was performed using *E. coli* and *S. aureus* AlexaFluor™488 BioParticles™ (Invitrogen), following manufacturer’s instructions. L3 wandering larvae carrying the *HmlΔdsred.nls* hemocytes marker were bled into 150 µL of Schneider’s insect medium (Sigma-Aldrich) containing 1 µM phenylthiourea (PTU, Sigma-Aldrich). The hemocyte suspension was then transferred to 1.5 mL low binding tubes (Eppendorf) and 2 x 10^7^ AlexaFluor™488 bacteria BioParticles™ were added. The samples were incubated at room temperature for 60 minutes to enable phagocytosis, and then placed on ice in order to stop the reaction. The fluorescence of extracellular particles was quenched by adding 0.4% trypan blue (Sigma-Aldrich) diluted 1/3. Phagocytosis was quantified using a flow cytometer (BD Accuri C6, USA) in order to measure the fraction of cells phagocytosing, and their fluorescent intensity. w^1118^ larvae and *HmlΔdsred.nls* larvae with or without bacterial particles were used to define the gates for hemocytes and the thresholds for phagocytosed particle emission. The phagocytic index was calculated as follows:

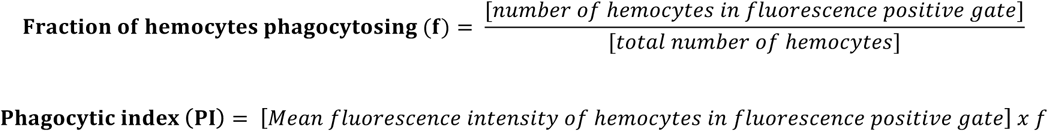
2. *Ex vivo* phagocytosis assays of *Serratia marcescens*, *Staphylococcus epidermidis*, and *Micrococcus luteus* was performed as follows. Bacterial strains and labelling bacteria with fluorescein isothiocyanate (FITC) are described in [50]. The stocks of *S. marcescens (*Szeged Microbial Collection, University of Szeged, Szeged, Hungary; SzMC 0567), *S. epidermidis (*SzMC 14531), and *M. luteus (*SzMC 0264) were used. Bacteria were conjugated by FITC as described by Zsámboki et al. Briefly, 10 ml of bacterial culture (OD_600_=1.5) was heat-inactivated in PBS and the cell pellet was re-suspended in 10 ml of 0.25 M carbonate-bicarbonate buffer pH 9.0. Fluorescein isothiocyanate (FITC) 0.5 mg, dissolved in 100 µl DMSO (Sigma) was added to the heat inactivated bacteria, rotated overnight at 4 °C and washed eight times with PBS. The FITC labelled bacteria were re-suspended, centrifuged at 11200 x g, the pellet was resuspended to a final concentration of 10%, sodium azide was added as a preservative (0.1%) and the samples were kept at 4°C till use. Bacteria were washed 5x with PBS prior to the phagocytosis assay. The phagocytic activity of hemocytes was assayed with a protocol similar to [6]. Hemocytes were isolated from third instar larvae at room temperature into Shields and Sang M3 insect medium (Sigma) containing 5% fetal-calf serum (Gibco) supplemented with 1-phenyl-2-thiourea (Sigma) to prevent melanization. A total of 2-3×10.5 hemocytes were incubated with 5-6 x 10^6^ heat-killed, FITC labeled bacteria at room temperature for 40 min in the wells of round bottomed microtiter plates (Gibco) in 100 µl. The fluorescence of extracellular bacteria was quenched by the addition of Trypan blue to the cells in 0.2% final concentration shortly before the actual measurement. The fluorescence intensity of phagocytosed FITC-labeled bacteria was analyzed with a FACS Calibur equipment (Beckton Dickinson). The data of three independent experiments were pooled. Phagocytic index was calculated as mentioned above.
3. Phagocytosis of green fluorescent 1 µm latex beads (Sigma-Aldrich) and AlexaFluor™488 Zymosan BioParticles™ (Invitrogen) was performed following the same procedure described above, with the exception that hemocytes without the *HmlΔdsred.nls* marker were used. 1 x 10^5^ Zymosan BioParticles™ and 0.2 µg of latex beads were added to each sample.

Given the difference in hemocyte numbers per larva between the genotypes, we bled 6 *w*^*1118*^ (*BL5905*) larvae, 4 *eater*^1^ and *NimC1*^1^ larvae, and 3 *NimC1^1^;eater^1^*larvae.

### Proliferation assays

Cell proliferation was assessed by 5-ethynyl-2′-deoxyuridine (EdU) labelling. Second instar, or middle L3 larvae were fed at 29°C with 1 mM 5-ethynyl-2 deoxyuridine (EdU) in fly food for 4 hours. Larvae were bled individually in 30 µL Schneider medium (Gibco) containing 1 nM phenylthiourea (PTU, Sigma). Hemocytes were allowed to settle for 30 minutes before being fixed in 4% paraformaldehyde PBS. Click-iT™ EdU Imaging Kit (Invitrogen) was used to stain *HmlΔdsred.nls* hemocyte populations. Cells were finally stained with 1/15000 dilution of 4’,6-diamidino-2-phenylindole DAPI (Sigma) and mounted in anti-fading agent Citifluor AF1 (Citifluor Ltd). The proliferation rate was determined by counting the number of EdU positive cells over the whole *HmlΔdsred.nls* hemocyte population. At least 6 animals were analysed per genotype.

### Scanning electron microscopy (SEM)

Samples for SEM were prepared as follows. Six wandering third instar larvae were bled into 50 µL of Schneider’s insect medium (Sigma-Aldrich) containing 1µM phenylthiourea (PTU, Sigma-Aldrich). The collected hemolymph was incubated on a glass coverslip for 20 minutes for spreading assay, or 30 minutes with bacteria for phagocytosis assay, before being fixed for one hour with 1.25% glutaraldehyde in 0.1 M phosphate buffer, pH 7.4. Samples were then washed in cacodylate buffer (0.1 M, pH 7.4), fixed again in 0.2% osmium tetroxide in the same washing buffer and then dehydrated in graded alcohol series. Samples underwent critical point drying and Au/Pd coating (4 nm). Scanning electron micrographs were taken with a field emission scanning electron microscope Merlin, Zeiss NTS, Germany.

### Transmission electron microscopy (TEM)

Third instar wandering larvae were bled in 50 µL of Schneider’s insect medium (Sigma-Aldrich) containing 1µM phenylthiourea (PTU, Sigma-Aldrich). The collected hemolymph was incubated with bacteria on a glass coverslip for one hour before being fixed for two hours with 2% paraformaldehyde + 2.5% glutaraldehyde in 0.1M phosphate buffer, pH 7.4. Samples were then washed in cacodylate buffer (0.1M, pH 7.4), fixed again in 1% osmium tetroxide and potassium ferrocyanide 1.5% in cacodylate buffer. After washes in distilled water, samples were stained in 1% uranyl acetate in water, washed again, and then dehydrated in graded alcohol series (2X50%, 1X70%, 1X90%, 1X95%, 2X100%). Embedding was performed first in 1:1 Hard EPON and ethanol 100%, and afterwards in pure EPON, before being embedded on coated glass slides and placed at 60°C overnight. Images were acquired with a FEI Tecnai Spirit 120 kV.

### Binding assay with live fluorescent bacteria

#### Cytochalasin D treatment

L3 wandering larvae were bled into 120 µL of Schneider’s insect medium (Sigma-Aldrich) containing 1 µM phenylthiourea (PTU, Sigma-Aldrich). Hemocytes were allowed to adhere on the glass slide for one hour before being treated for another 60 minutes with 1 µM of Cytochalasin D. After drug treatment, hemocytes were incubated directly on the slide with live fluorescent *S. aureus* or *E. coli* bacteria always in the presence of Cytochalasin D for 60 minutes. After fixation in 4% paraformaldehyde PBS, rhodamine phalloidin staining (Molecular Probes™) was performed. Finally, cells were stained with 1/5000 dilution of 4’,6-diamidino-2-phenylindole DAPI (Sigma) and mounted in anti-fading agent Citifluor AF1 (Citifluor Ltd.).

#### Phagocytosis inhibition by cold temperature

L3 wandering larvae were bled into cold Schneider’s insect medium (Sigma-Aldrich) containing 1 µM phenylthiourea (PTU, Sigma-Aldrich) on a previously chilled glass slide. After larva bleeding, hemocytes and bacteria were incubated directly on the pre-chilled slide, in cold Schneider’s medium, on ice for 60 minutes. Fixation and staining procedures were performed as described above.

#### Fluorescent bacteria

The *E. coli*-GFP strain was obtained by transforming *E. coli* K12 with a synthetic sfGFP coding sequence cloned in a pBAD backbone (ThermoFisher) by Gibson assembly. sfGFP induction was obtained by growing the bacteria in LB + 0.1% arabinose overnight prior to the binding assay. The *S. aureus*-GFP strain is described in [59]

### Hemocyte phalloidin staining and cell area measurement

Five wandering third instar larvae were bled on a microscope slide into 120 µL of 1X PBS containing 1 µM phenylthiourea (PTU, Sigma-Aldrich). The hemocytes were then allowed to adhere for 30 minutes, before being fixed in PBS, 4% paraformaldehyde. Phalloidin staining was performed with diluted 1/100 AlexaFluor488-or rhodamine-phalloidin (Molecular Probes™). Finally, cells were stained with a 1/5000 dilution of 4’,6-diamidino-2-phenylindole DAPI (Sigma) and mounted in anti-fading agent Citifluor AF1 (Citifluor Ltd.). Samples were imaged with an Axioplot Imager.Z1 Zeiss coupled to an AxioCam MRm camera (Zeiss).

For cell area measurements, hemocytes were captured with a 20x objective on GFP, RFP and DAPI channels. Individual images were then loaded into a CellProfiler pipeline (www.cellprofiler.org). In order to define the cell area, cell nuclei were first detected using data from the DAPI channel. Cell limits were then defined by expanding the nuclei signal to the edges of the GFP channel. Cell areas were computed from this segmentation analysis, and cell area of 750 cells of each genotype was quantified.

### Crystal cell counting methods

At least ten third instar larvae were heated in 1 mL of phosphate-buffered saline (PBS) at 67°C for 20 minutes in ppendorf tubes. For quantification analysis, black puncta were counted in the posterior-most segments A6, A7 and A8. Pictures were taken with a Leica DFC300FX camera and Leica Application Suite right after heating.

For quantification of crystal cells by flow cytometry, we crossed *w*^*1118*^ or mutant *lzGal4,UAS-GFP* flies with the corresponding *HmlΔdsred.nls* wild-type or mutant flies. Larvae form the resulting offspring were used to determine the number of crystal cells (*lzGal4,UAS-GFP*) and the ratio of crystal cells among the total hemocyte population (*lzGal4,UAS-GFP* / *HmlΔdsred.nls*). Four larvae of each genotype were bled into 150 µL 1X PBS containing 1µM phenylthiourea (PTU, Sigma-Aldrich) and 0.1% paraformaldehyde to block crystal cell rupture. 75 µL of the hemocyte suspension was analysed by flow cytometry. Hemocytes were first selected from debris by plotting FSC-A against SSC-A on a logarithmic scale in a dot plot. Cells were then gated for singlets by plotting FSC-H versus FSC-A. FL1 and FL2 detectors were used for *lzGal4,UAS-GFP* and *HmlΔdsred.nls* events, respectively.

### Wounding experiment

Wandering third-instar larvae were pricked dorsally near the posterior end of the animal, using a sterile needle (diameter ∼5 µm). Pictures of melanised larvae were taken 20 minutes after pricking, with a Leica DFC300FX camera and Leica Application Suite.

### Wasp Infestation and quantification of fly survival to *L. boulardi* wasp infestation

For wasp infestations experiments, 30 synchronized second instar (L2) larvae were placed on a pea-sized mound of fly food within a custom-built wasp trap in the presence of three female*L. boulardi* wasps for 2 hours. Quantification of fly survival was performed as follows. Parasitized larvae were kept at room temperature and scored daily for flies or wasps emergence. The difference between enclosed flies and wasps to the initial number of larvae was set as dead larvae/pupae. Pictures of melanised eggs were taken with a Leica DFC300FX camera and Leica Application Suite 70 hours after wasp infestation.

### Infection experiments and qRT-PCR

Systemic infections (septic injuries) were performed by pricking third instar larvae dorsally near the posterior end of the animal using a thin needle previously dipped into a concentrated pellet (OD_600_ ∼200) of bacteria. After septic injury, larvae were incubated at 29°C. After 4 hours, the animals were collected, and total RNA extraction was performed using TRIzol reagent (Invitrogen). RNA quality and quantity were determined using a NanoDrop ND-1000 spectrophotometer and 500 ng of total RNA was used to generate cDNA using SuperScript II (Invitrogen, Carlsbad, California, United States). Quantitative PCR was performed on cDNA samples using the LightCycler 480 SYBR Green Master Mix (Roche). Expression values were normalized to *Rp49*.

### Statistical analysis

Experiments were repeated at least three times independently and values are represented as the mean ± standard deviation. Data were analysed using GraphPad Prism 7.0. *p*-values were determined with Mann-Whitney tests. * *p*<0.05, ** *p*<0.01, *** *p*<0.001, ns: not significant.

## Results

### Generation of a *NimC1* null mutant by homologous recombination

In order to characterize *NimC1* functions, we generated a null mutant by deleting the corresponding *NimC1* gene region. The deletion removes the ATG translation start site and the following 852 bp sequence. The knockout was performed in the *w*^*1118*^ background, using homologous recombination [33], which also lead to the insertion of a 7.9 kb cassette carrying the *white*^*+*^ gene (Fig S1A-B). Functional deletion of *NimC1* was confirmed by RT-PCR performed on total RNA (Fig S1C). As *NimC*1 is specifically expressed in hemocytes and has been implicated in phagocytosis, we combined the *NimC1* mutation with the previously described *eater*^1^ null mutant [30], generating a double mutant *NimC1^1^;eater^1^*. Both *NimC1*^1^ and *NimC1^1^;eater^1^* flies were viable and did not show any developmental defect. For over-expression studies, we also generated flies containing the *NimC1* gene downstream of the *UAS* promoter. Using these tools, we characterized the function of NimC1 focussing on hemocytes of third instar larvae.

### *NimC1* deficient hemocytes show adhesion defects *in vitro*

Eater has been involved in hemocyte adhesion and sessility [30]. Given the structural similarities between NimC1 and Eater [27], we first investigated the role of NimC1 in cell adhesion. We observed that the cell area of *NimC1*^1^ and *eater*^1^ adherent hemocytes was decreased compared to that of *w*^*1118*^ wild-type control (Fig 1A). Notably, the cell area of *NimC1^1^;eater^1^* adherent hemocytes was significantly smaller than that of single mutants. Quantification analysis revealed that wild-type hemocytes have a mean cell area of 237 µm^2^, while *NimC1*^1^*, eater*^1^ and *NimC1^1^;eater^1^* mutants have 120, 114 and 99.7 µm^2^, respectively (Fig 1B). Image-based cytometry analysis of free-floating hemocytes revealed that the spreading defects observed in our mutants were not due to an inherently smaller cell size (Fig 1C). Of note, over-expression of *NimC1*, using the *HmlΔ,Gal4* plasmatocyte driver, did not increase the adhesion properties of the hemocytes (Fig S2A-C).

**FIGURE 1.**
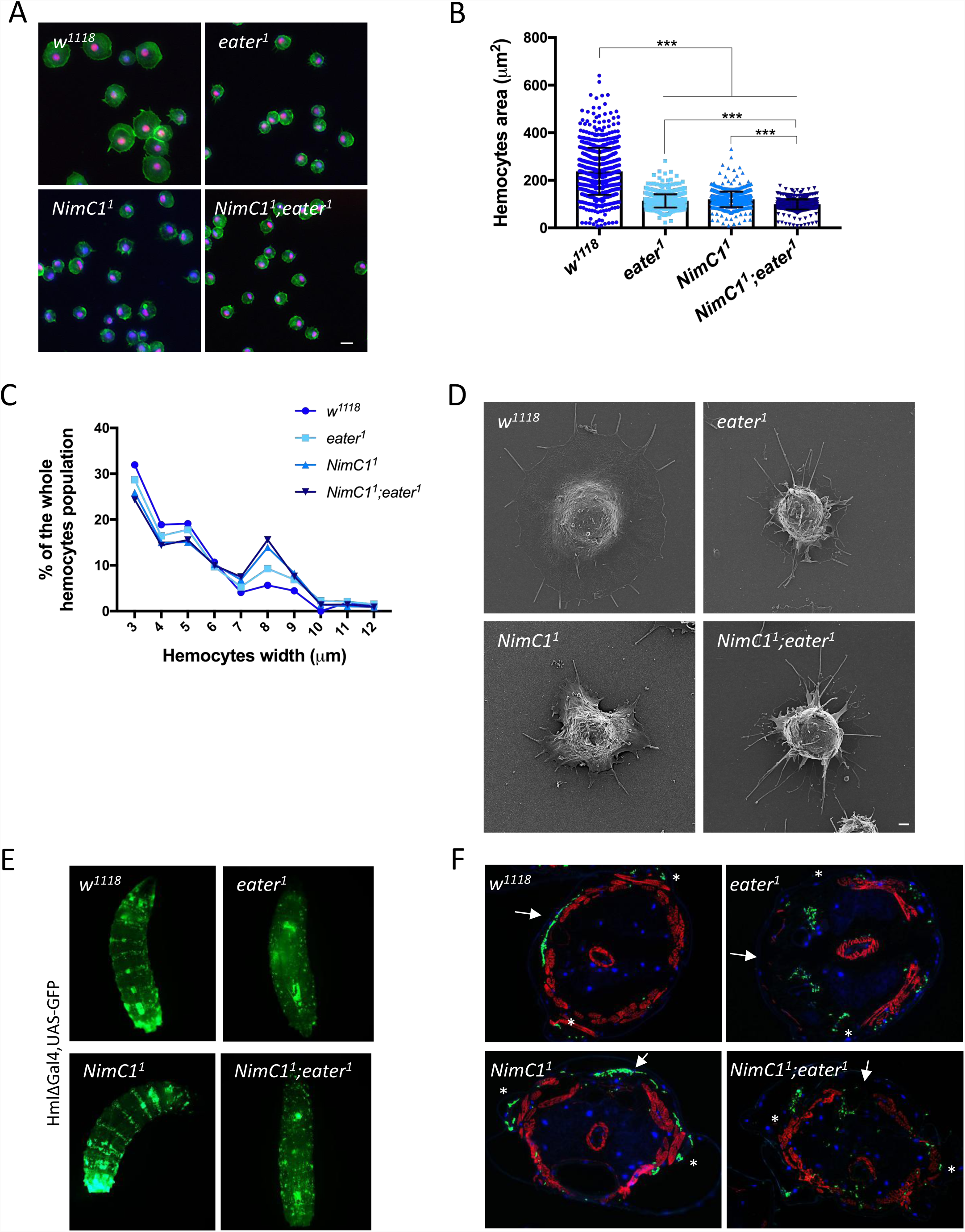
*NimC1*^1^ hemocytes show spreading defects *in vitro*. **(A)** Representative images of fixed hemocytes from *w*^*1118*^, *eater*^1^, *NimC1*^1^ and *NimC1^1^;eater^1^* L3 wandering larvae combined with *Hml*^*Δ*^*dsred.nls* marker (red). Hemocytes of the indicated genotypes were extracted by larval bleeding, allowed to spread for 30 minutes on a glass slide, and stained with AlexaFluor™488 phalloidin (green). Scale bar: 10 µm. (**B)** Mean cell area quantification of fixed *HmlΔdsred.nls* hemocytes spread for 30 minutes on slides and stained with AlexaFluor488 phalloidin. Cell area of 750 cells was quantified using the CellProfiler software. (**C)** Size distribution of free floating hemocytes from *w*^*1118*^, *eater*^1^, *NimC1*^1^ and *NimC1^1^;eater^1^* L3 wandering larvae. Hemocyte size was measured with TALI imaged-based cytometer directly after larval bleeding. (**D)** Scanning electron microscopy images of spread hemocytes from the indicated genotypes of L3 wandering larvae. Scale bar: 1 µm. (**E)** Whole larva images of *w*^*1118*^, *eater*^1^, *NimC1*^1^ and *NimC1^1^;eater^1^* third instar larvae specifically expressing *UAS-GFP* in plasmatocytes driven by *HmlΔ,GAL4.* The dorsal side of the animal is shown. (**F)** Cross sections of the indicated genotype from L3 wandering larvae combined with *HmlΔGal4,UAS-GFP (*green). Rhodamine phalloidin staining (red) was performed after larva cross sectioning. Cell nuclei are shown in DAPI (blue). White arrows and asterisk indicate dorsal and lateral side of the animal, respectively.

In order to get a better insight into these adhesion defects, we investigated lamellipodia and filopodia morphology by scanning electron microscopy (SEM) on spread hemocytes. Lamellipodia are a key feature of highly motile cells, playing a central role in cell movement and migration [34]. They represent flat cellular protrusion, characterized by an enriched network of branched actin filaments. Filopodia, instead, are rather used by the cell to sense the surrounding microenvironment, and consist of parallel actin filaments that emerge from the lamellipodium. Plasmatocytes from wild-type larvae appeared as round adherent cells with a central bulge within the cell body, from which lamellipodia and filopodia extended. *NimC1* and *eater* null hemocytes were still able to form narrow filopodia projections. However, *NimC1* hemocytes showed a smaller lamellipodia region, which was much more strongly reduced in *eater* hemocytes. Furthermore, *NimC1;eater* double mutant hemocytes exhibited a phenotype similar to that of *eater (*Fig 1D). Collectively, our results point to a role of NimC1 in hemocyte spreading.

Previous work has shown that *eater* larvae lack the sessile hemocyte compartment and have all peripheral hemocytes in circulation [30]. To further investigate whether the *NimC1* deletion affects sessility, we explored hemocyte localization using the hemocyte marker *HmlΔ,Gal4>UAS-GFP* by whole larva imaging and cross section visualization. In *NimC1*^1^*,HmlΔ>Gal4,UAS-GFP* third instar (L3) wandering larvae, hemocytes were still able to enter the sessile state, forming dorsal and lateral patches (Fig 1E-F). In contrast, both *eater*^1^ and *NimC1^1^;eater^1^* larvae lacked sessile hemocytes, all plasmatocytes being in circulation (Fig 1E-F). Finally, over-expression of *NimC1* in hemocytes from *eater* deficient larvae did not rescue the lack of sessility phenotype and the *ex vivo* adhesion defect caused by the absence of *eater (*Fig S2D). *In vivo* RNAi targeting *NimC1* confirmed the hemocyte adhesion defect observed with the null mutant (Fig S3A-B). This indicates that the observed phenotypes were indeed caused by the deletion of *NimC1* and not the genetic background. Altogether, our data indicate that NimC1 contributes to hemocyte adhesion *ex vivo*, but in contrast to Eater, it is not directly required for hemocyte sessility *in vivo*.

### *NimC1* null larvae have an increased number of hemocytes

In order to further investigate the role of NimC1 and its potential interaction with Eater in peripheral haematopoiesis, we counted by flow cytometry the number of sessile and circulating hemocyte populations. Third instar larvae containing the hemocyte marker *HmlΔdsred.nls*, combined with the *NimC1* and *eater* null mutants, were used. Our study confirmed that *eater* L3 wandering mutant larvae have more hemocytes than the wild type [30] (Fig 2A). Similarly, *NimC1*^1^ third instar larvae have 3.2 times more circulating hemocytes compared to the wild-type (Fig 2A). As *NimC1*^1^ L2 larvae have a wild-type like number of hemocytes, the increase in hemocyte counts in this mutant takes place at the end of larval development (Fig 2B). Surprisingly, hemocyte numbers were 6 times higher in *NimC1^1^;eater^1^* double mutant larvae (Fig. 2A), suggesting that *eater* and *NimC1* additively regulate hemocyte counts. A higher hemocyte number was already observed in second instar larvae in the double mutant (Fig 2B). Dissection of the lymph gland, revealed no obvious differences in size between mutants and wild type control (Fig 2C). We then explored whether the increased hemocyte count observed in our mutants was caused by a higher proliferation rate. EdU incorporation experiments revealed that *NimC1*^1^ and *eater*^1^ single mutants have a higher frequency of peripheral proliferating hemocytes compared to wild-type in middle L3 but not L2 larvae (Fig 2D-E). The higher proliferation rates might therefore explain the increased number in hemocyte counts in both middle L3 (Fig 2F) and L3 wandering (Fig. 2A) larvae. Interestingly, we found that both hemocyte count and mitotic rate were higher in *NimC1^1^;eater^1^* in L2 and L3 larvae indicating that both receptors additively regulate hemocyte proliferation levels (Fig 2A-B, D-F). The higher hemocyte count in *NimC1* mutant larvae was phenocopied when using the *in vivo* RNAi approach (Fig S3C).

**FIGURE 2.**
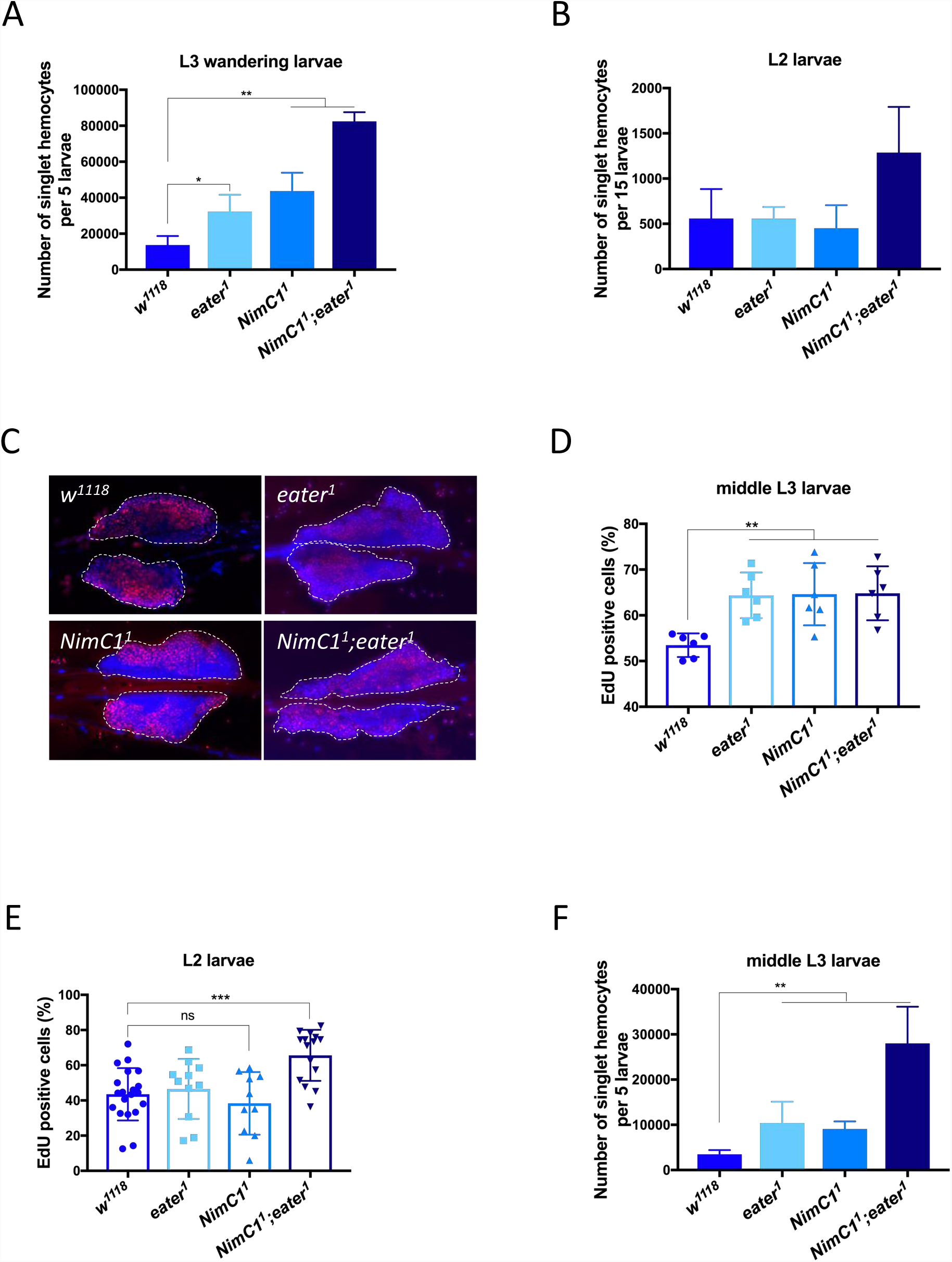
*NimC1*^1^ and *NimC1^1^;eater^1^* larvae give rise to a higher number of hemocytes. Number of singlet peripheral hemocytes per 5 L3 wandering (**A)** and 15 L2 (**B)** larvae of *w*^*1118*^, *eater*^1^, *NimC1*^1^, *NimC1^1^;eater^1^* combined with *HmlΔdsred.nls*. (**C)** Lymph glands of *w*^*1118*^, *eater*^1^, *NimC1*^1^, *NimC1^1^;eater^1^* mutant third instar larvae combined with *HmlΔdsred.nls* hemocyte marker (red). Lymph gland primary lobes are shown and boundaries are delimited by a white dashed line. Cell nuclei were stained with DAPI (blue). (**D-E)** Percentage of EdU positive cells upon *HmlΔdsred.nls* cells in middle L3 (D) and L2 larval (E) stage of *w*^*1118*^, *eater*^1^, *NimC1*^1^, *NimC1^1^;eater^1^*. A number of at least 6 animals were used for each genotype.**(F)** Number of singlet peripheral hemocytes per 5 middle L3 larvae of the indicated genotypes, combined with *HmlΔdsred.nls*.

We also investigated a possible role of NimC1 in crystal cell and lamellocyte differentiation. Crystal cells are the second hemocyte type present in non-infected larvae, specifically involved in the melanization response and wound healing. Crystal cells can be found in both the sessile and circulating state. Recent studies have shown that a fraction of those cells derive from sessile plasmatocyte by transdifferentiation [8], [23]. Consequently, it has also been shown that crystal cells need sessile plasmatocytes to be themselves sessile [30]. Lamellocytes are barely present in healthy larvae, but can differentiate from plasmatocytes [21], [35] or prohemocytes [36] in response to specific stress signals, such as parasitization. They are thought to play an essential role in encapsulation of parasitoid wasp eggs. Our data indicate that *NimC1* mutants retain the ability to differentiate fully mature crystal cells and lamellocytes (Fig S4). Finally, we did not uncover any role of NimC1 in the systemic antimicrobial response of larvae against Gram-positive (*Micrococcus luteus*) or Gram-negative bacteria (*Erwinia carotovora carotovora*), as revealed by the wild-type like induction of *Diptericin* and *Drosomycin* gene expression, two target genes of the Imd and Toll pathways, respectively [37] (Fig S5).

### NimC1 contributes with Eater to phagocytosis of bacteria

A previous *in vivo* RNAi approach had revealed a role of NimC1 in the phagocytosis of Gram-positive bacteria [27]. We used the *NimC1* deletion to further elucidate the requirement of this receptor in bacterial uptake by performing *ex vivo* phagocytosis assays. As previously reported [30], *eater* null mutant hemocytes were impaired in their capacity to phagocytose the Gram-positive bacterium *S. aureus (*Fig 3A), but not the Gram-negative bacterium *E. coli (*Fig 3B). In contrast to the previous RNAi experiments [27], loss of *NimC1* affected neither the phagocytosis of *S. aureus* nor that of *E. coli (*Fig 3A-B). However, hemocytes derived from *NimC1;eater* mutant larvae were not only severely impaired in the phagocytosis of *S. aureus (*Fig 3A), as expected, but also of *E. coli (*Fig 3B). This indicates that Eater and NimC1 contribute redundantly to the phagocytosis of Gram-negative bacteria, as the presence of Eater or NimC1 is able to compensate for the absence of the other. The use of a double mutant also revealed a contribution of NimC1 to the phagocytosis of *S. aureus*, although Eater plays the predominant role. To further confirm these phagocytosis defects, we extended the analysis to two additional Gram-positive (Fig 3C-D) (*Staphylococcus epidermidis*, *Micrococcus luteus*) and one Gram-negative (Fig 3E) (*Serratia marcescens*) bacteria. In agreement with our previous results, *NimC1*^1^ hemocytes did not show phagocytosis defects for *S. epidermidis*, *M. luteus* and *S. marcescens* FITC-labeled bacteria (Fig 3C-E). However, phagocytosis of all those microbes was strongly impaired in *NimC1^1^;eater^1^* double mutant hemocytes, further confirming our initial findings (Fig 3C-E).

**FIGURE 3.**
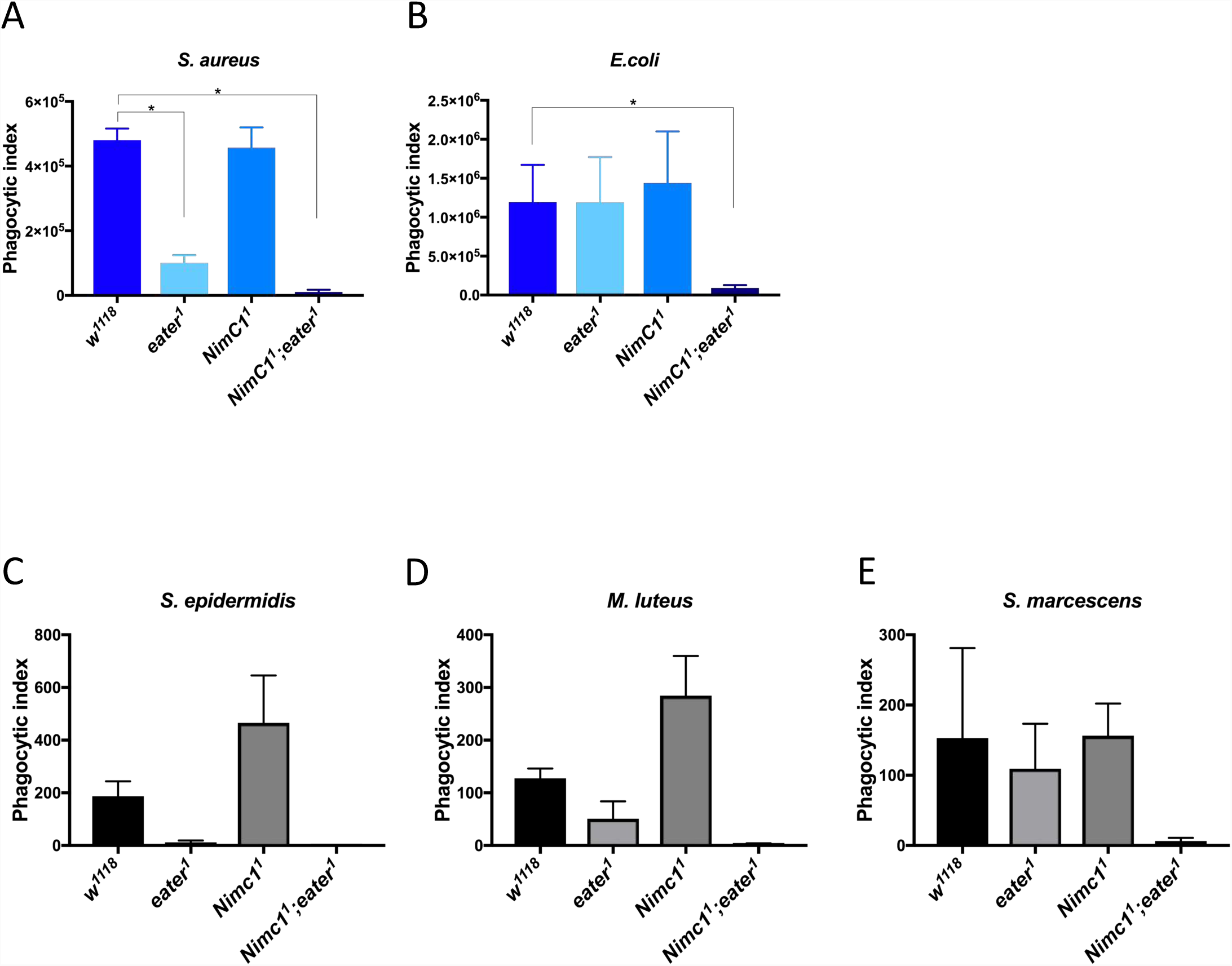
NimC1 contributes with Eater to phagocytosis of Gram-positive and Gram-negative bacteria. **(A-B)** *Ex vivo* phagocytosis assay using *Staphylococcus aureus (*A) and *Escherichia coli (*B) AlexaFluor™488 BioParticles™ (Invitrogen). *HmlΔdsred.nls* hemocytes from third instar wandering larvae were incubated with the particles for 60 minutes at room temperature. (**C-E)** *Ex vivo* phagocytosis assay using the Gram-positive *Staphylococcus epidermidis (*C) and *Micrococcus luteus (*D) and Gram-negative *Serratia marcescens (*E) bacteria. Bacteria were first heat-inactivated and subsequently labelled with fluorescein isothiocynate (FITC). Hemocytes from third instar wandering larvae of the indicated genotypes were incubated with the bacteria for 40 minutes at room temperature. The data of three independent experiments are shown. In A-D, phagocytosis was quantified by flow cytometry, and the fluorescence of extracellular particles quenched by adding trypan blue.

### Eater and NimC1 receptors play a critical role in adhesion to bacteria

To better understand the cause of *eater*^1^ and *NimC1*^1^*,eater*^1^ phagocytosis defects, and thereby to elucidate the unique role of these receptors in bacterial uptake, we performed scanning (SEM) and transmission (TEM) electron microscopy experiments. Both these techniques allow following the different membrane-driven events during the phagocytosis process. Hemocytes from the corresponding genotypes were incubated with either *E. coli* or *S. aureus* live bacteria for 30 minutes to evaluate bacterial adhesion by SEM, and to follow bacterial uptake at 60 minutes by TEM. In wild-type and *NimC1*^1^ hemocytes incubated with *S. aureus*, we observed plasma membrane remodelling, with the formation of a phagocytic cup and pseudopod protrusions that progressively surrounded bacteria, finally leading to their engulfment (Fig 4A-B, white arrowheads). Similar observations were made for *wild-type*, *NimC1*^1^, and *eater*^1^ hemocytes incubated with *E. coli* bacteria (Fig 4C-D). Surprisingly, upon incubation of *eater*^1^ and *NimC1^1^;eater^1^* hemocytes with *S. aureus*, no bacteria were present on the cell surface (Fig 4A). A decreased level of bacteria adherence was also observed in *NimC1^1^;eater^1^* hemocytes incubated with *E. coli (*Fig 4C). In accordance with SEM experiments, transmitted electron micrographs showed numerous engulfment events in *wild-type* and *NimC1*^1^ hemocytes with *S. aureus* bacteria (Fig 4D, arrows), as well as for *E. coli* in *wild-type*, *NimC1*^1^, and *eater*^1^ hemocytes. Altogether, these experiments point to the importance of Eater in binding Gram-positive bacteria, which is consistent with a previous report [38], but also to a redundant role of NimC1 and Eater in binding Gram-negative bacteria. Furthermore, they suggest that these two receptors do not play any critical role in bacteria internalization, as all three mutants showed (rare) engulfment events.

**FIGURE 4.**
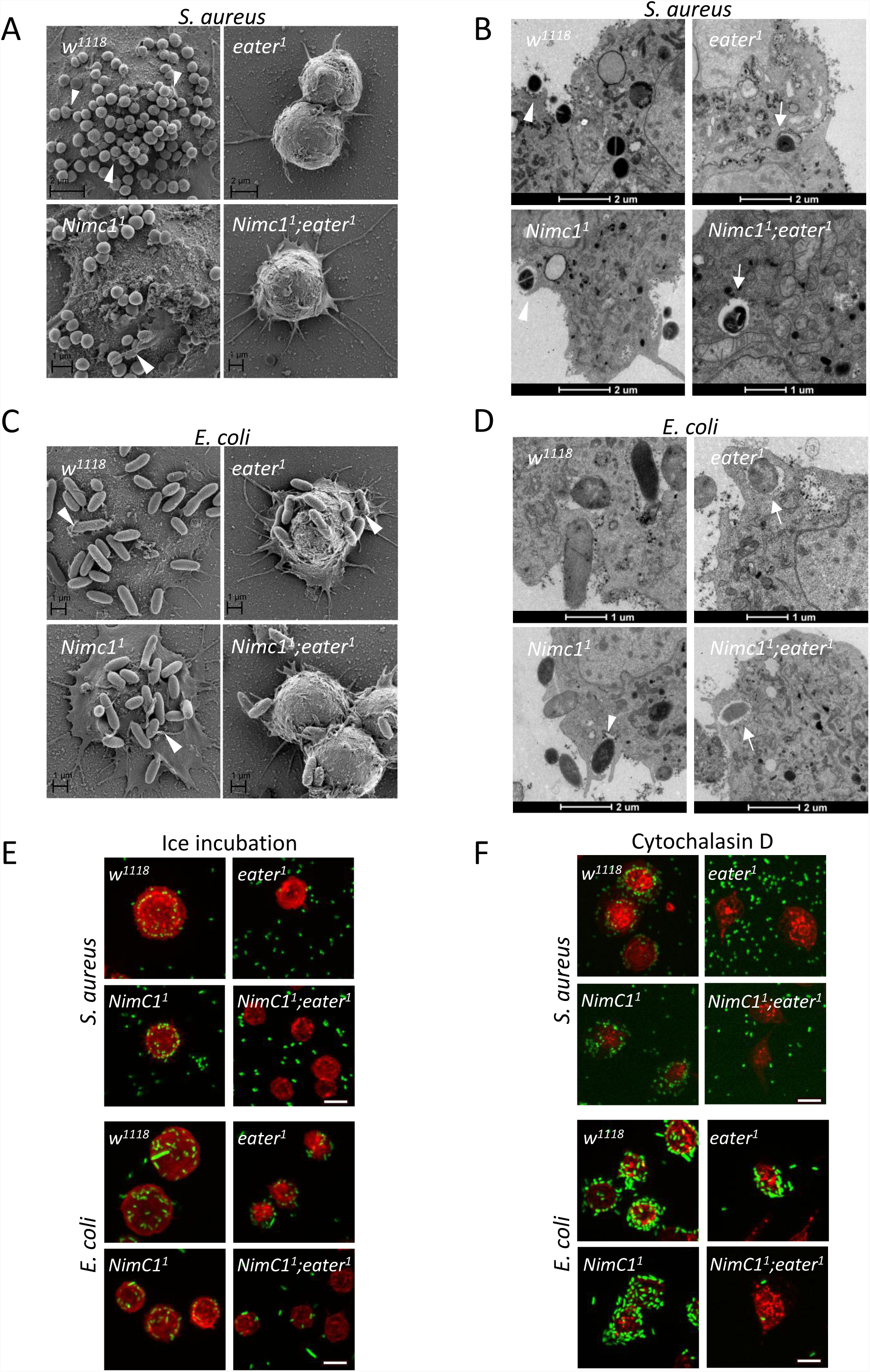
*eater*^1^ and *NimC1^1^;eater^1^* hemocytes show bacteria adhesion defects. **(A)** Scanning electron microscopy images of hemocytes from the indicated genotypes of L3 wandering larvae after 30 minutes incubation with *Staphylococcus aureus* live bacteria, at room temperature. (**B)** Transmission electron micrographs of hemocytes from the indicated genotypes of L3 wandering larvae after 60 minutes incubation with *Staphylococcus aureus* live bacteria, at room temperature. (**C)** Scanning electron microscopy images of hemocytes from the indicated genotypes of L3 wandering larvae after 30 minutes incubation with *Escherichia coli* live bacteria, at room temperature. (**D)** Transmission electron micrographs of hemocytes from the indicated genotypes of L3 wandering larvae after 60 minutes incubation with *Escherichia coli* live bacteria, at room temperature. Arrowheads and arrows indicate hemocyte membrane protrusions and internalized bacteria, respectively. (**E-F)** Hemocytes from the corresponding genotypes were incubated with *Staphylococcus aureus* GFP (*green*) or *Escherichia coli* GFP (*green*) live bacteria on ice (E), or with Cytochalasin D (F), for one hour (see Material and Methods section for further details). After fixation with 4% paraformaldehyde, hemocytes were stained with rhodamine phalloidin (*red*). Scale bar: 10 µm.

To further confirm the bacteria adhesion defects, we incubated hemocytes and live fluorescent bacteria either on ice or with Cytochalasin D. Both treatments inhibit the engulfment process, without altering the binding of the bacteria to the phagocytic cell [6]. In both conditions (Fig 4E-F), we did observe less bacteria binding to plasmatocytes in the same genotypes that were defective for phagocytosis in our *ex vivo* assays (*eater*^1^ for *S.* aureus, and *NimC1^1^;eater^1^* for *S. aureus* and *E. coli*, Fig 3A-B). The drastic effect observed with the *NimC1^1^;eater^1^* double mutant on phagocytosis and bacteria adhesion led us to explore the contribution of other previously characterized phagocytic receptors using the same assays. Draper and Croquemort are two transmembrane receptors expressed by plasmatocytes, and belong to the Nimrod and CD36 family, respectively [27], [39]. With SIMU (NimC4), they both play a key role in the engulfment of apoptotic bodies [32], [39]–[42]. Moreover, a role in *S. aureus* phagocytosis has also been described for Draper and Croquemort, as well as in *E. coli* phagocytosis for Draper [31], [43]–[45]. Although we observed a modest decrease in *S. aureus* phagocytosis in *croquemort* and *draper* mutants (called *crq*^*Δ*^ and *drpr*^*Δ^5^*^ respectively), and *E. coli* in *drpr*^*Δ^5^*^ (Fig S6A-B), bacteria adhesion to the hemocytes was not impaired in these mutants (Fig S6C). This further supports a specific role of Eater as the main tethering receptor in Gram-positive bacteria phagocytosis.

### Phagocytosis of Latex Beads and Zymosan yeast particles is impaired in *NimC1* null mutants

To further understand the role of Eater and NimC1 in the phagocytosis process, we proceeded to analyse the uptake of “neutral” latex beads particles. We also tested their role in the phagocytosis of zymosan, a compound found on the cell wall of yeast. While bacteria present at their surface specific targets for the engulfing receptors, latex beads can be seen as non-immunogenic particles, that do not bear any ligands for the phagocyte. We observed that the phagocytic index of latex beads was wild type-like in *eater* null plasmatocytes, as well as in *drpr*^*Δ^5^*^ and *crq*^*Δ*^ (Fig 5A, Fig S6D). This further indicates a specific role of Eater in microbe uptake, likely via the recognition of a key bacterial surface determinant. Interestingly, plasmatocytes lacking the NimC1 receptor showed a significantly reduced ability to engulf latex beads, as well as zymosan yeast particles (Fig 5A-B). Thus, we could uncover a phagocytic defect in the *NimC*1 single mutant only when using particles that do no display bacterial motifs, suggesting that bacteria can bypass NimC1, probably by recruiting other phagocytic receptors, such as Eater.

**FIGURE 5.**
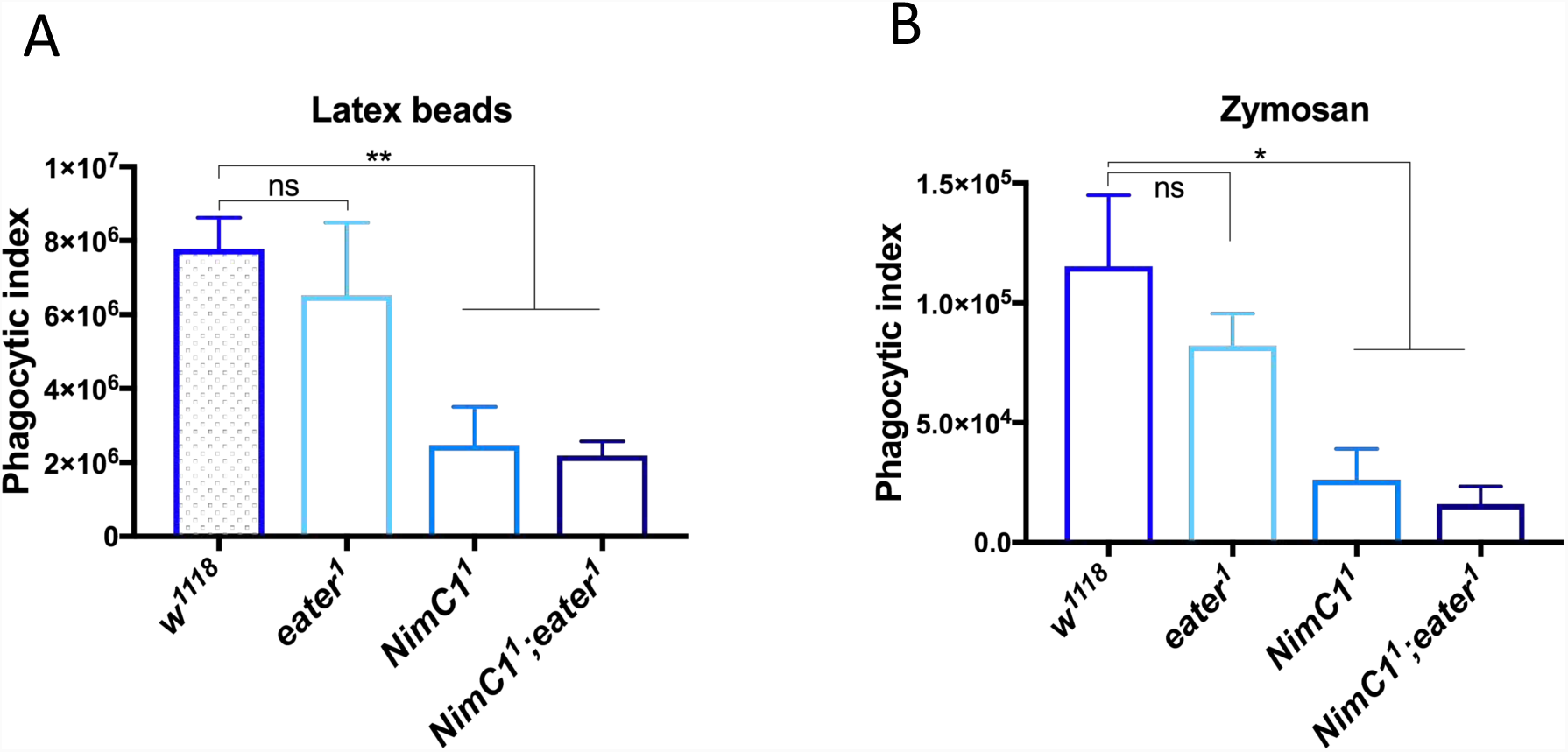
*NimC1*^1^ hemocytes show impaired phagocytosis of latex beads and zymosan particles. **(A)** Phagocytic index quantification of latex beads (Sigma-Aldrich) engulfment in hemocytes from *w*^*1118*^, *eater*^1^, *NimC1*^1^ and *NimC1^1^;eater^1^* L3 wandering larvae after 30 minutes incubation. (**B)** Phagocytic index quantification of AlexaFluor™488 Zymosan (*S. cerevisiae*) BioParticles™ engulfment. Zymosan BioParticles were incubated with *w*^*1118*^ or mutant hemocytes from L3 wandering larvae for 90 minutes. In (A) and (B), phagocytosis was quantified by flow cytometry, and the fluorescence of extracellular particles quenched by adding trypan blue.

## Discussion and Conclusions

NimC1 was initially identified as an antigen for the plasmatocyte specific monoclonal antibody P1. It belongs to the Nimrod gene family that has been implicated in the cellular innate immune response in *Drosophila* [46], [47]. A previous study pointed to the importance of NimC1 in the phagocytosis of bacteria, since RNAi-mediated silencing of this gene resulted in decreased *S. aureus* uptake by plasmatocytes [27]. In the present work, we further re-evaluated the function of the NimC1 protein by using a novel null-mutant, revealing its precise role in hemocyte adhesion, proliferation and phagocytic ability.

By performing *ex vivo* spreading assays, we observed that the cell area of adherent *NimC1* null hemocytes was reduced compared to wild-type control, suggesting that NimC1 works as an adhesion molecule. Consistent with this observation, scanning electron microscopy on spread hemocytes of *NimC1*^1^ mutants revealed a defect in lamellipodia extension. Spreading defects were also observed in hemocytes lacking Eater [30]. Thus, two structurally related Nimrod receptors, NimC1 and Eater, are involved in hemocyte adhesion. Interestingly, the electron micrographs revealed diverse spreading phenotypes between *NimC1* and *eater* knockout adherent plasmatocytes. According to these experiments, Eater appeared to play a more important role in lamellipodia extension, since this defect was less marked in *NimC*^1^ hemocytes. It will be interesting to analyse, in future work, the implications of NimC1 and Eater in hemocyte migration during metamorphosis or wound healing. Our results also indicate that Eater and NimC1 additively regulate hemocyte adherence. In contrast to *eater* deficient larvae, NimC1 is however not directly required for plasmatocyte sessility *in vivo*. Whether NimC1 contributes to hemocyte sessility through additional scaffold proteins has to be further investigated, even though the present evidence might favour a model where Eater is the only essential protein required for hemocyte sessility [30].

During larval development, the peripheral hemocyte population undergoes a significant proliferation, expanding by self-renewal [8], [22]. Moreover, during these developmental stages, plasmatocytes are characterized by a dynamic behaviour, continuously exchanging between the sessile and circulating state. In 2011, Makhijani et al. provided evidence that plasmatocyte proliferation rate is higher in the hematopoietic pockets, where hemocytes cluster on the lateral side of the larval body [22]. At this location, sessile plasmatocytes are in contact with the endings of peripheral neurons, which are thought to provide a trophic environment to the blood cells. More recently, it has been shown that sensory neurons of the peripheral nervous system produce Activin-ß, which turned out to be an important factor in the regulation of hemocyte proliferation and adhesion [48]. By analysing the total number of hemocytes in third instar *NimC1*^1^ or *eater*^1^ larvae, we observed that both Eater and NimC1 negatively regulate hemocyte counts in an additive manner. EdU incorporation experiments revealed that the higher hemocyte counts in *NimC1^1^;eater^1^* mutants were a consequence of an increased hemocyte proliferation rate. It is tempting to speculate that the higher proliferation rate is a secondary consequence of an adhesion defect. Indeed, adherent cells, notably when establishing contacts with other cells, are less proliferative, a process called “contact inhibition of proliferation” [49]. Future studies should address how Eater and NimC1 contribute to both adhesion and proliferation, and the direction of causality between these two processes remains to be disentangled. Like plasmatocytes, crystal cells increase in number during larval stages. However, crystal cell proliferation is not due to a self-renewal mechanism because mature crystal cells do not divide. Instead, a recent study has shown that new crystal cells originate from transdifferentiation of sessile plasmatocytes via a Notch-Serrate dependent process [23]. In the present study, we show that the *NimC1* deletion does not strongly impact crystal cell formation as both sessile and circulating crystal cell populations were only mildly affected in *NimC1* null larvae. Moreover, NimC1 does not affect the ability to encapsulate parasitoid wasps. These data indicate that our mutants retain the ability to fully differentiate mature crystal cells and lamellocytes.

NimC1 was initially identified as a phagocytic receptor, mediating the uptake of *S. aureus* bacteria [27]. Contrary to this study, our *ex vivo* phagocytosis assays using the *NimC1* deletion mutant revealed that the uptake of both Gram-positive and Gram-negative bacteria was not altered in *NimC1* null hemocytes. We hypothesized that the RNAi approach could have targeted other phagocytic receptors, revealing a stronger phenotype not observed in the single null mutant. Strikingly, phagocytosis of both bacteria types was severely impaired in *NimC1^1^;eater^1^* hemocytes, suggesting that both receptors contribute synergistically to phagocytosis of both Gram-negative and Gram-positive bacteria. At this stage, we cannot exclude that these receptors might indirectly regulate phagocytosis by controlling another receptor directly involved in bacterial recognition, although we judge this hypothesis unlikely. Consistent with our hypothesis, an RNAseq analysis of *eater* deficient versus wild-type hemocytes did not uncover any role of Eater in the regulation of other phagocytic receptors (data not shown). Given the marked phagocytosis defect of the *eater* single mutant against *S. aureus*, the contribution of NimC1 was especially noticeable in the case of the Gram-negative bacterium *E. coli*. Our scanning electron microscopy approach revealed that NimC1 and Eater might contribute together to the early step of bacterial recognition, since *NimC1^1^;eater^1^* double mutants showed decreased bacterial adhesion. The involvement of NimC1 in *E. coli* binding is consistent with previous *in vitro* work showing that native NimC1 binds bacteria [50]. Surprisingly, *NimC1*^1^ and *NimC1^1^;eater^1^*, but not *eater* deficient plasmatocytes, showed a significantly reduced ability to engulf latex beads and yeast zymosan particles. Thus, the two receptors have specific properties with regards to phagocytosis. It is interesting to address a parallel with the implication of two Nimrod receptors in bacteria tethering and docking, as shown for apoptotic cells clearance in *Drosophila melanogaster* [32], [51]. Tethering receptors usually lack an intracellular domain and are involved in the binding to the dying cell. Docking receptors, instead, are subsequently required to activate intracellular signalling and mediate the internalization and degradation of the particle. In the fruit fly, a good example for tethering and docking receptors are SIMU/NimC4 and Draper, respectively [51]. A similar dichotomy exists in vertebrates, as Stabilin 2 and TIM-4 are classified as tethering receptors, whereas the integrins αVβ3 and αVβ5 are grouped as docking/signalling proteins [52], [53]. The involvement and cooperation of two receptors of the Nimrod family in bacterial phagocytosis raised the possibility that they might contribute via different mechanisms: binding and internalization. Our current hypothesis is that Eater might work as the main tethering receptor, required for binding to specific motifs present on the bacterial surface. Moreover, given the wild-type engulfment of latex beads in *eater*, this receptor might be engaged specifically for phagocytosis of microbes. Indeed, the involvement of Eater in bacterial binding was already assessed in previous studies [38], and is consistent with our assays using live fluorescent bacteria and SEM experiments. On the contrary, NimC1 could function in the activation of the subsequent intracellular signalling, maybe as a subunit of a bigger macromolecular complex. We hypothesise that in the presence of cell wall bacterial determinants, such as peptidoglycan, lipopolysaccharide or teichoic acids, microbe phagocytosis can bypass the requirement of NimC1 by providing enough “eat me” signals to Eater. In contrast, the critical role of NimC1 in phagocytosis becomes visible with less immunogenic particles. This would explain why we do not observe any defects in the phagocytosis of *S. aureus* and *E. coli* in *NimC1* single mutant, but only with latex beads (i.e. particles without any bacterial motifs).

Future studies should address how Eater and NimC1 interact, the implication of other possible phagocytic receptors and characterization of their respective ligands. Collectively, our study identifies NimC1 and Eater as two critical receptors involved in the initial step of phagocytosis, which might be the best window to directly recognize bacterial ‘eat me’ signals that trigger phagocytosis. While a plethora of receptors have been identified for their role in microbe phagocytosis, our genetic analysis using compound mutants unambiguously reveals the major role and contribution of Eater and NimC1 to this process. Our study also provides a valuable tool to better assess the role of phagocytosis during the immune response.

## Authors contributions

CM and BL conceived and designed the project. CM, AJB, and ER contributed to the generation of the *NimC1*^1^ mutant and other tools used in this study. JD performed the wasp experiments, and EK and IA did the phagocytosis assay with *S. epidermidis*, *M. luteus*, and *S. marcescens*. CM performed all the other experiments of the study. CM and BL wrote the paper.

## Acknowledgments

We thank Jean-Paul Vincent, Cyril Alexandre and Alberto Baena for gene targeting tools and Samuel Rommelaere and Claudine Neyen for critical reading of the manuscript. We thank the BioEM platform (EPFL) for their important help with the electron microscopy experiments, the Flow Cytometry Core Facility (EPFL) for their constant assistance in the use of the Accuri DB machine, and the BIOP platform (EPFL) for their help using confocal microscopes and designing the Cell pipeline analyser. We would like to thank Florent Masson for the *E. coli GFP* strain used in this study, Jean-Philippe Boquete for his precious technical support, and Mickael Poidevin for construction of plasmids.

## Supplementary Figure Legends

**FIGURE S1.**
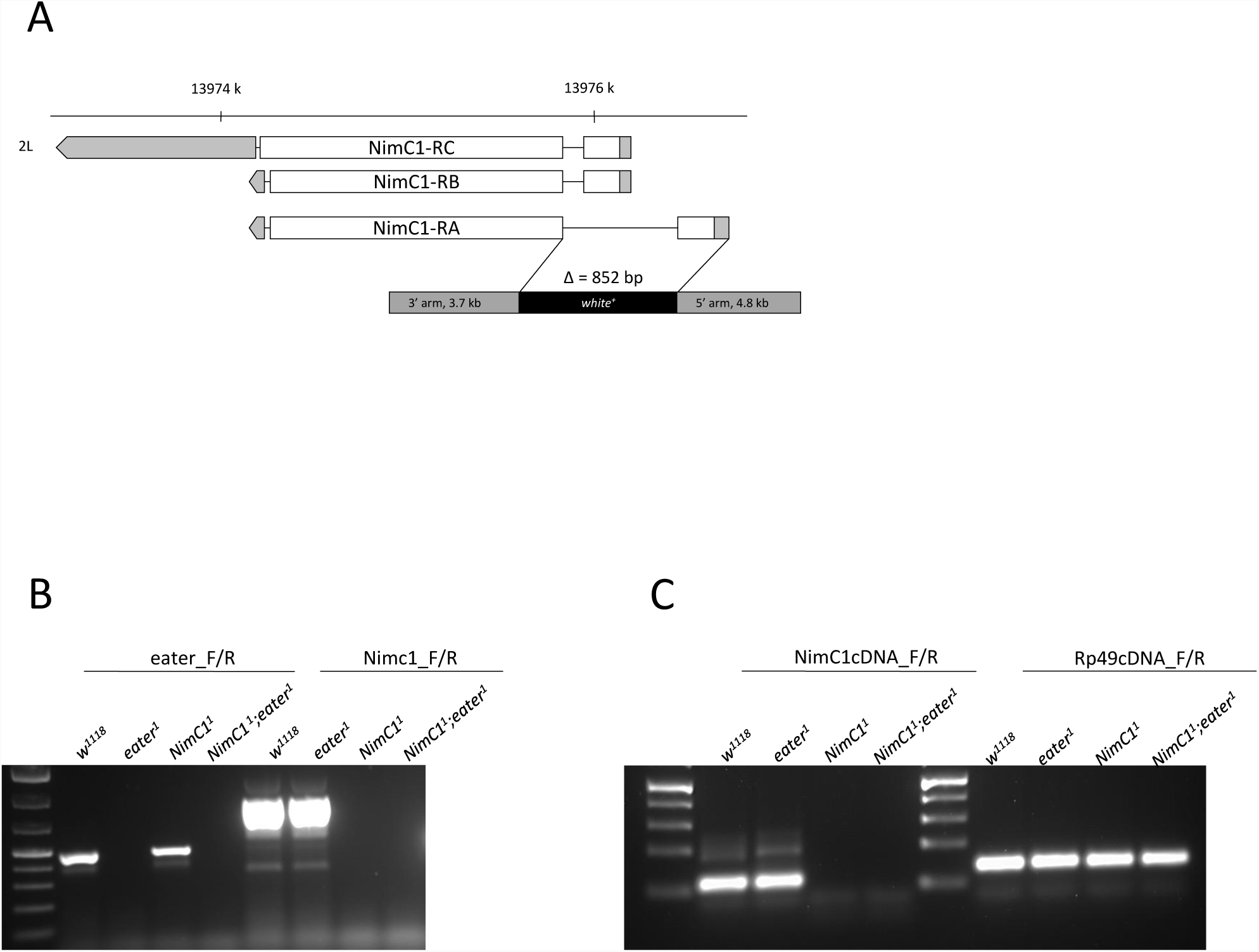
Gene targeting and deletion of *NimC1*. **(A)** *NimC1* gene deletion by homologous recombination. The *NimC1* gene is located on the left (L) arm of chromosome 2 and it encodes three isoforms. White and grey boxes represent exons and UTR regions, respectively. Eye colour was transformed from white to red by the *white*^+^ marker. (**B)** PCR genotyping confirming the targeted deletion of the *NimC1* gene, whereas the *eater* locus was not affected. (**C)** RT-PCRs confirming functional deletion of *NimC1*.

**FIGURE S2.**
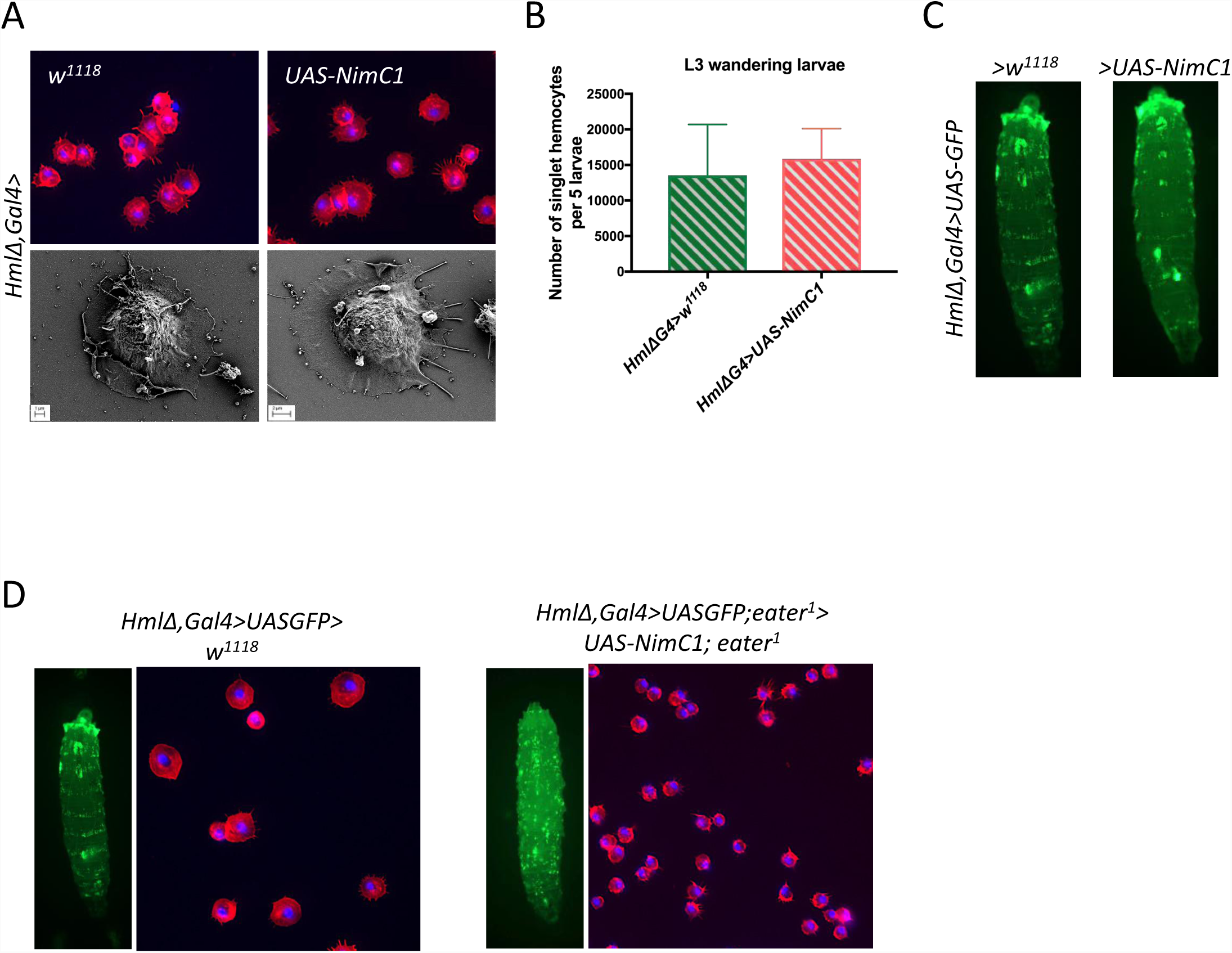
*NimC1* overexpression does not alter hemocyte adhesive properties. **(A)** Over-expression of *NimC1* does not alter the spreading properties of hemocytes. Upper panel: Representative images for fixed hemocytes from *HmlΔ,Gal4*>*w*^*1118*^ and *HmlΔGal4*>*UAS-NimC1* L3 wandering larvae stained with rhodamine phalloidin (red). Cell nuclei are shown in DAPI (blue). Bottom panel: scanning electron micrographs on spread hemocytes from *HmlΔ,Gal4*>*w*^*1118*^ and *HmlΔGal4*>*UAS-NimC1* of L3 wandering larvae. (**B)** Number of singlet peripheral hemocytes in third instar wandering larvae is not affected upon NimC1 overexpression. Results are represented as a sum of 5 animals with the indicated genotypes. (**C)** Whole larva imaging of *HmlΔGal4,UAS-GFP>w*^*1118*^ and *HmlΔGal4,UAS-GFP>UAS-NimC1* shows no major difference in hemocyte localization pattern when NimC1 is specifically overexpressed in hemocytes. The dorsal side of the animal is shown. (**D)** Whole larva imaging and spreading assay showing the absence of rescue when overexpressing the *NimC1* in an *eater* mutant background.

**FIGURE S3.**
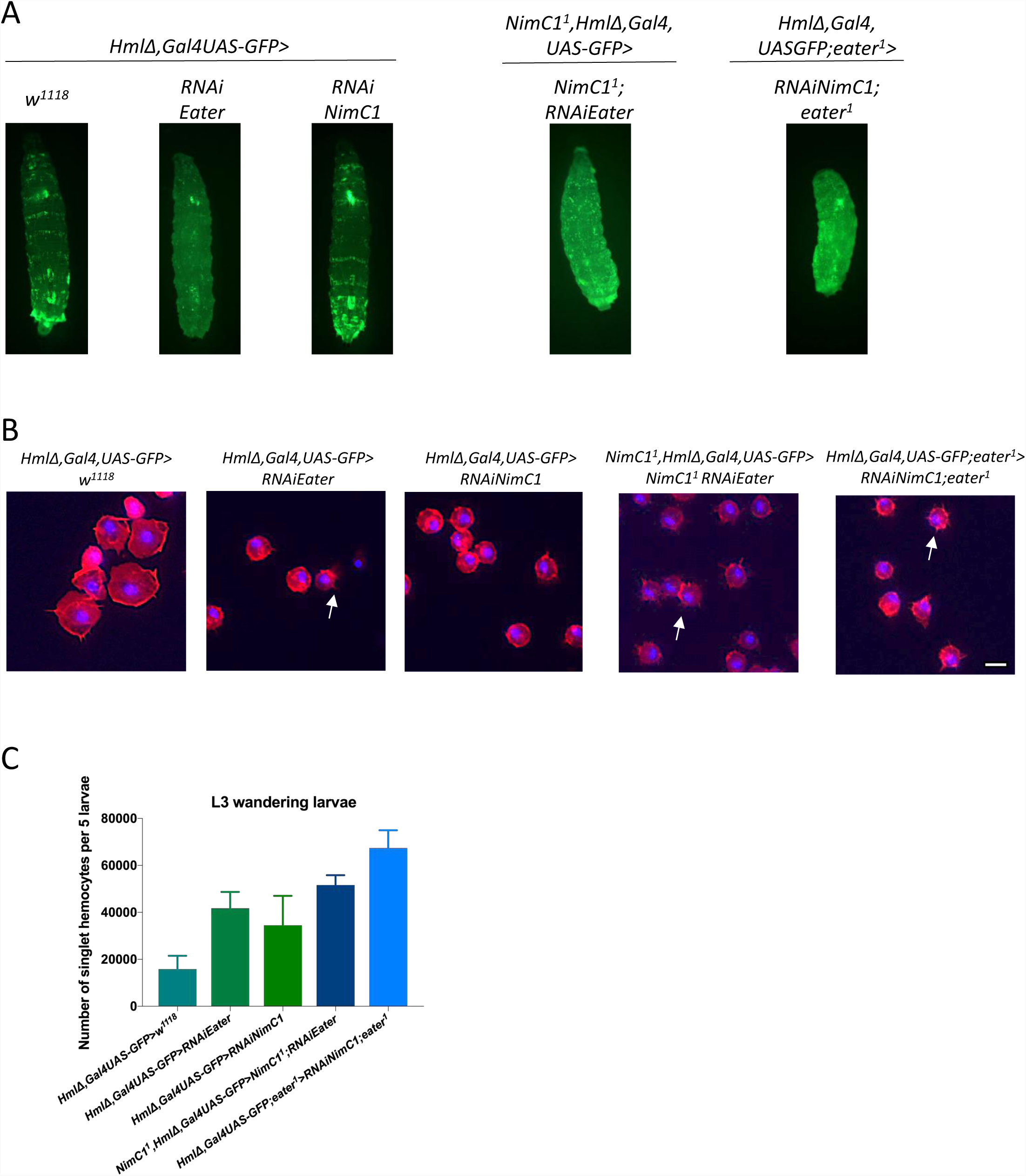
RNAi mediated knockdown of *NimC1* and *eater* confirmed *NimC1*^1^ and *NimC1^1^;eater^1^* phenotypes. **(A)** Whole larva images of third instarlarvae of the indicated genotypes, specifically expressing GFP in plasmatocytes driven by *HmlΔ,GAL4*. The dorsal side of the animal is shown. (**B)** Representative images of fixed hemocytes from the indicated genotypes of L3 wandering larvae, stained with rhodamine phalloidin (red). Cell nuclei were stained with DAPI (blue). Arrows indicate the presence of filopodia in the corresponding genotype, found also in the *eater* deletion mutants. Scale bar: 10 µm. (**C)** Number of singlet peripheral hemocytes per 5 L3 wandering larvae of the indicated genotypes.

**Figure S4.**
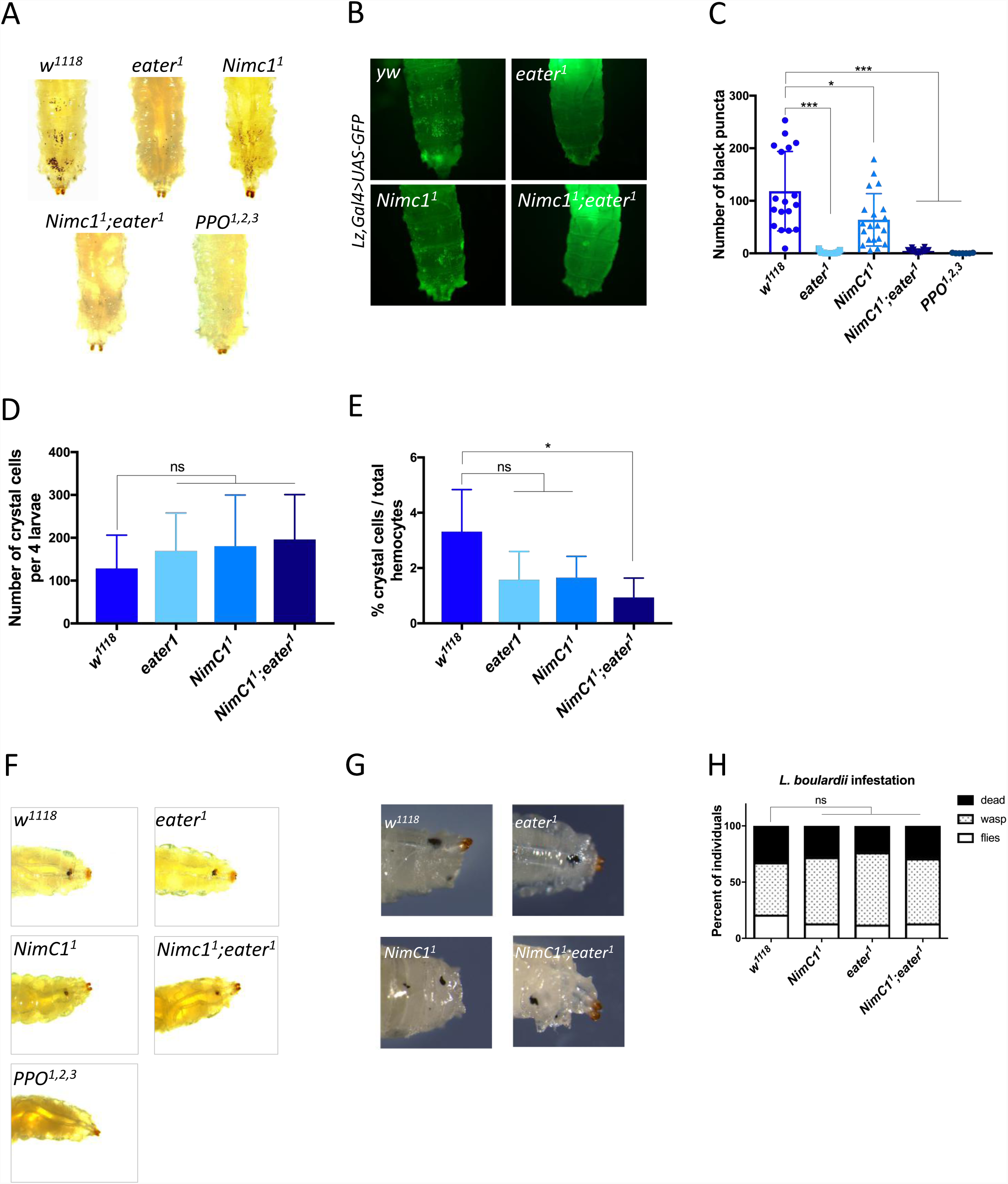
Sessile and circulating crystal cell populations are mildly affected in *NimC1^1^* larvae. (**A**) Heating larvae induces the spontaneous activation of the prophenoloxidase zymogen within crystal cells, leading to their blackening [54]. Consequently, the population of crystal cells attached under the cuticle becomes visible as black puncta. Sessile crystal cell numbers were mildly reduced in *NimC1*^1^ larvae, while being almost completely absent in *eater*^1^ and *NimC1^1^;eater^1^* larvae. Shown are representative images of *w*^*1118*^, *eater*^1^, *NimC1*^1^ and *NimC1^1^;eater^1^* L3 wandering larvae after heat treatment at 67°C for 20 minutes. (**B)** *In-vivo* imaging using the *lzGal4,UAS-GFP* crystal cell marker confirmed the previous observations. Shown is a dorsal view of the five posterior-most segments in *yw*, *eater*^1^, *NimC1*^1^ and *NimC1^1^;eater^1^* L3 wandering larvae, previously combined with the crystal cell lineage marker *lzGal4,UAS-GFP*. (**C)** Black puncta count from the three posterior-most segments of heated *w*^*1118*^*, eater*^1^*, NimC1*^1^ *and NimC1*^1^*;eater*^1^ third instar larvae. (**D-E)** Counting of *lzGal4,UAS-GFP* positive cells revealed a wild-type number of crystal cells in *NimC1* and *eater* deficient L3 wandering larvae, and a moderately decreased ratio of crystal cells over the total hemocyte population, pointing to a mild defect in sessility but not in the general ability to differentiate crystal cells. (D) Number of singlet *lzGal4,UAS-GFP* labelled crystal cells per 4 third instar wild-type or mutants larvae. (E) Ratio of *lzGal4,UAS-GFP* upon *HmlΔdsred.nls* positive cells in L3 wandering larvae of the indicated genotypes. Crystal cells counting in D and E was performed by flow cytometer analysis. (**F**) Melanization response to epithelial wounding is wild-type like in Eater and NimC1 mutants larvae. Images of melanized larvae were acquired 20 minutes after pricking. (**G-H)** We investigated the ability of *NimC1* mutant larvae to encapsulate wasp eggs upon parasitization. For this, we studied larvae upon infestation with the parasitoid wasp *Leptopilina boulardi*, which induces a massive production of lamellocytes. (G) All mutant lines showed a wild-type ratio of melanotic encapsulation of *L. boulardi* wasp eggs after infestation. Shown are representative images of melanised wasp egg in *w*^*1118*^ control and mutants, 70 hours after *L. boulardi* infestation. (H) Quantification of emerging *D. melanogaster* adult, *L. boulardi* wasp, and dead animals. Data are shown as a sum of three experiments, with a total of 90 animals for each genotype. Data were analysed using Chi-square statistical test (*p*-value > 0.05). In (A) and (F) PPO^1^,^2^,^3^ mutant larvae were used as negative control.

**FIGURE S5.**
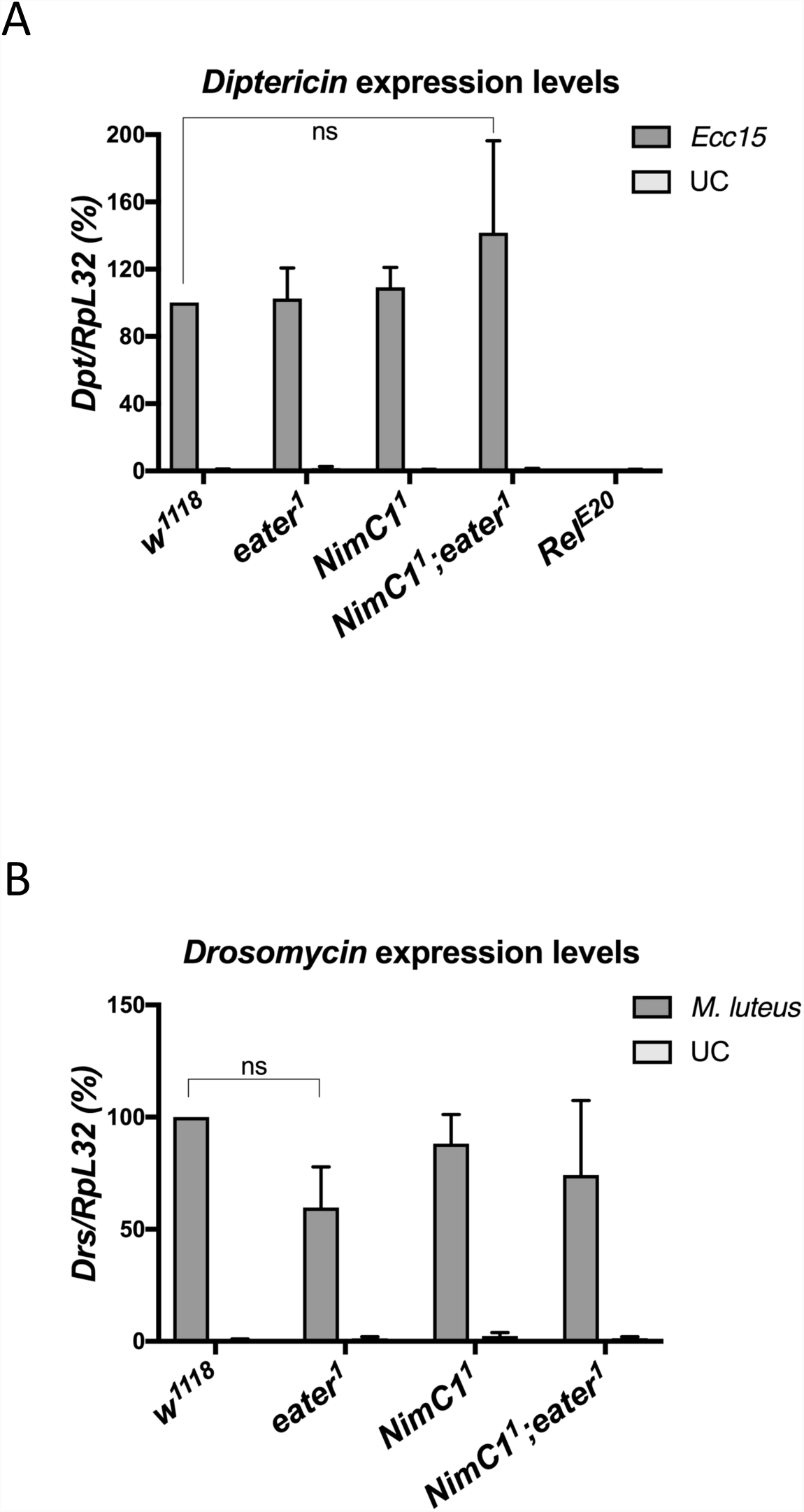
NimC1 is not involved in humoral immunity. Expression of *Diptericin (***A)** and *Drosomycin (***B)** in *w*^*1118*^, *eater*^1^, *NimC1*^1^ and *NimC1^1^;eater^1^* third instar larvae. Total RNA from infected animals was extracted 4 hours after *Erwinia carotovora carotovora (Ecc15)* or *Micrococcus luteus (M. luteus)* septic injury. UC= unchallenged controls. Expression levels of *Diptericin (Dpt*) (A) and *Drosomycin (Drs*) (B) relative to *RpL32*. The Imd pathway mutant *Relish* was used as an immune-deficient control in (A).

**FIGURE S6.**
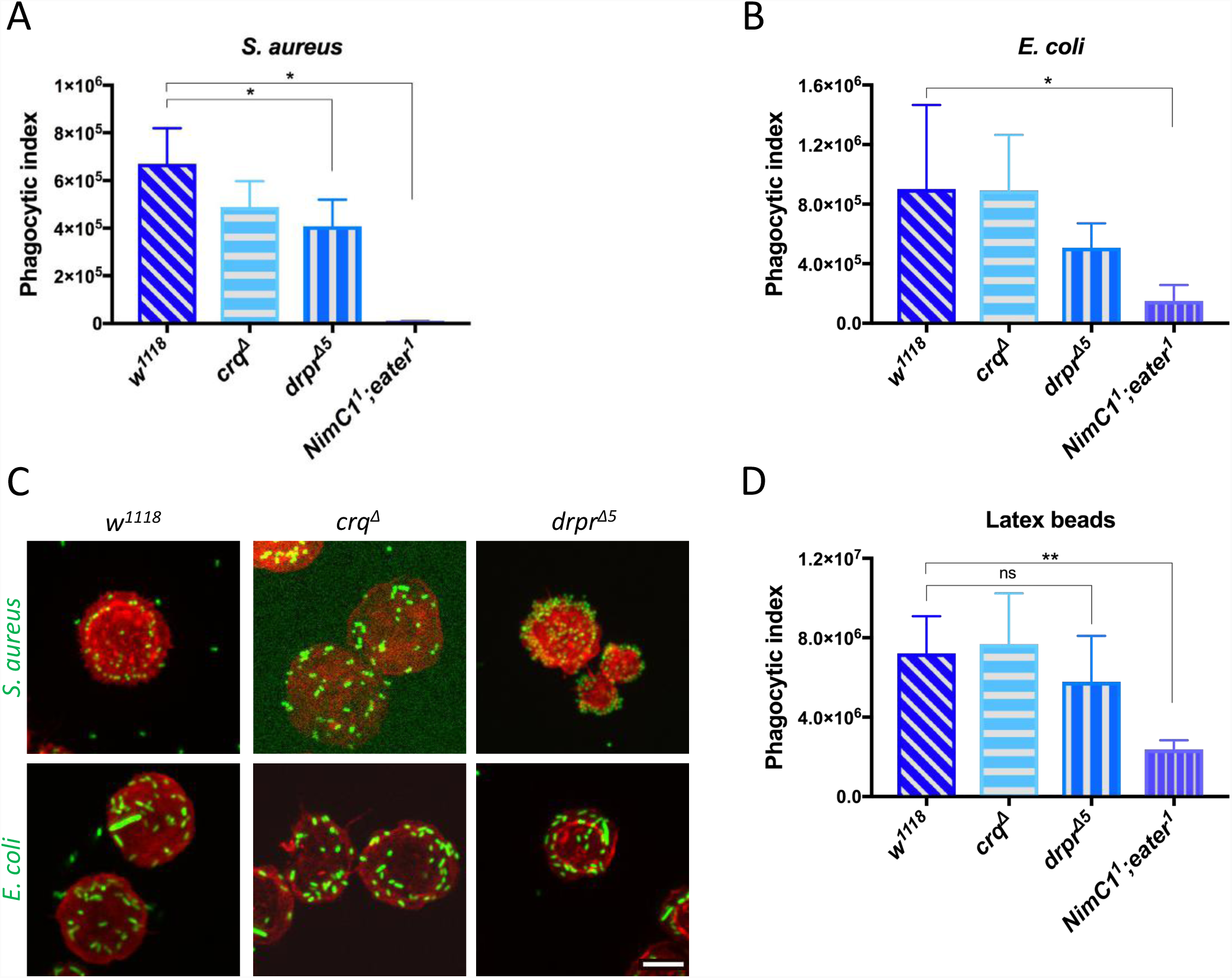
Phagocytosis in *draper* and *croquemort* mutants. Phagocytosis of *S. aureus (***A)** and *E. coli (***B)** AlexaFluor™488 BioParticles™ (Invitrogen) and latex beads (Sigma-Aldrich) (**C)** in *crq*^*Δ*^ and *drpr*^*Δ^5^*^ hemocytes mutants from L3 wandering larvae. *w*^*1118*^ and *NimC1^1^;eater^1^* hemocytes were used as positive and negative controls, respectively. (**E)** *crq*^*Δ*^ and *drpr*^*Δ^5^*^ hemocytes mutants show no binding defect of *S. aureus (*upper panel) and *E. coli (*bottom panel) bacteria. Hemocytes from the corresponding genotypes were incubated with live GFP bacteria (green) on ice for one hour (see Material and Methods section for further details). After fixation with 4% paraformaldehyde, hemocytes were stained with rhodamine phalloidin (red). Scale bar: 10 µm.

